# Seed Selection Strategies for Overlap Detection

**DOI:** 10.1101/316273

**Authors:** Jonathan Teutenberg

## Abstract

The current state-of-the-art assemblers of long, error-prone reads rely on detecting all-vs-all overlaps within the set of reads with overlaps represented by a sparse selection of short subsequences or “seeds”. Though the quality of selection of these seeds can impact both accuracy and speed of overlap detection, existing algorithms do little more than ignore over-represented seeds. Here we propose several more informed seed selection strategies to improve precision and recall of overlaps. These strategies are evaluated against real long-read data sets with a range of fixed seed sizes. We show that these strategies substantially improve the utility of individual seeds over uninformed selection.

## 1 Background

Recent advances in sequencing technologies such as those of Pacific Biosciences and Oxford Nanopore Technologies have enabled the generation of reads that are orders of magnitude longer but also more error-prone than those produced by high quality short read technologies. These long reads – potentially up to two million bases in length – are currently best assembled by detecting overlaps between read pairs[3, 4]. Example algorithms for overlap detection include MHAP[1] (as used by the Canu assembler), minimap2[8] (as used by the mini-asm assembler[7]), the TULIP assembler[5], and BLAT[6].

A common feature of these overlap detection implementations is the representation of reads as a sparse collection of short subsequences or *seeds*. Overlap detection is then performed as an all-vs-all comparison amongst these collections, based on exact matches of seeds. The selection of the seeds used to represent a read should be sparse (so as to reduce computational complexity), shared by a maximal number of other overlapping reads (to increase true positives), and shared by a minimal number of non-overlapping reads (to decrease false positives).

Sparsity may be ensured either by selecting non-overlapping seeds at defined intervals (e.g. BLAT), by selecting a ‘minimal value’ seed within a given window (e.g. minimap2), or by selecting a fixed number of seeds that each minimise their own value function (e.g. MHAP). While all of these perform some form of seed selection, alternatives do exist. For example DALIGNER [9] maintains all seeds of a given length though when comparing a pair of reads it makes use of only the intersection of their collections of seeds.

In addition to sparsity, these algorithms also aim to minimise the number of overlaps in which their seeds appear, typically by increasing the seed length *k* and ignoring seeds that appear in the data with unusually high frequency.

The third criteria – maximising the number of overlapping reads containing a seed – is ignored by these algorithms. For example, minimap2 uses a lexicographic ordering for the hash function to determine the best ‘minimal’ local seed; MHAP uses a different random hash function for selecting each seed; and BLAT makes an arbitrary selection of equally spaced seeds.

One simple method to increase the proportion of overlapping reads containing a seed is to reduce *k*, though this becomes a trade-off with minimising its presence in non-overlapping reads. Perhaps the closest existing approach to addressing this is TULIP which selects seeds from contigs generated by highly accurate short reads[5]. These kinds of accurate seeds have been shown to greatly improve the resulting assemblies[2].

Selection from accurate short read data is a binary filter only - doesn’t rank seeds. Anything eliminated is guaranteed false positive which is nice. Better for larger k as fewer errors conflate with good seeds.

In this paper we consider alternative methods for ranking seeds that are expected to improve the recall of overlapping reads in comparison to the lexicographic ordering used currently.

## 2 Selection strategies

The term *selection strategy* is used here to describe a mapping

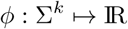
from the set of all possible subsequences length *k* (over, for example, an alphabet Σ of four nucleotide bases) to a value. During seed selection, an algorithm is then expected to use this strategy to compare k-mers, selecting that which maps to the maximal *ϕ* value when all other criteria of the algorithm are met.

A k-mer *u* is said to be *correct* in a read *s* (or its collection of k-mers *S*, that we shall use interchangeably below) when its bases match those at the same position in the sequence being read, written *c*(*u, s*). Otherwise it is *incorrect*.

An *overlap* at k-mer *u* in a read *s*_0_ is a set of reads that contain a k-mer at the same position in the sequence they have been read from. When an overlap detection algorithm must choose between two nearby k-mers *a* and *b*, we wish to select the one that is expected to appear in the greatest number of other sequences in the overlap *O*. Our aim is therefore to find a strategy for which

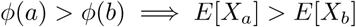
with *X_u_* the random variable describing the number |{*S_i_*|*u* ∊ *S_i_* ∀ *S_i_* ∊*O*}| of sequences containing the seed *u*. As seed presense is a binary value we have *E*[*X_u_*] = *np* where *p* is probability of *u* being in a k-mer collection *S_i_*, which we shall write *p_u_*. As *n* is fixed for all seeds in the overlap, it can be divided out and we now require that

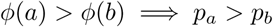

That is, given k-mers *a* and *b* in a sequence, if the selection strategy prefers *a*, then *a* is more likely to be in an overlapping sequence than *b*.

Note that in two overlapping sequences (i.e. that share a reference sequence), a k-mer *a* found in both is either correct in both or incorrect in both. So we have

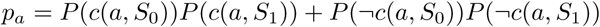
where *S*_1_ is any sequence in *O*. However, given a relatively low error rate and errors distributed amongst many possible incorrect k-mers, the probability of *a* being incorrect in both is insignificant in comparison to it being correct in both, therefore

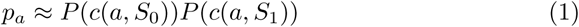
where the probability of being correct *P* (*c*(*a, S*_1_)) is the same for all *S_i_* ∊ *O*.

Later we will consider alternatives, but for our initial seed selection strategies we assume that a k-mers likelihood of being correct is dependent only on its base content i.e. it is independent of the context or read it is found in. Thus *P*(*c*(*a, S_i_*)) is a constant *P*(*c*(*a*)) for all *S_i_* containing *a*, and in addition *P*(*c*(*a, S*_0_) shares the same probability. Therefore,

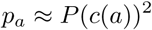
This assumption means an ideal selection strategy *ϕ* is one that maintains the same ordering of k-mers as their probability of being correct in the sequences in which they appear.

### 2.1 Quality scores

An obvious source of information on the probability that a potential seed is correct is the quality score provided by the base-caller that generated the read. In terms of the quality of a seed, we take the quality value assigned to the centre base (i.e. at offset *k/*2 from the seed’s position in the read). Other reasonable alternatives are to use the minimum quality within the seed, or perhaps the mean quality value.

As a strategy we define *ϕ_q_*(*a*) to be the mean quality value of all instances of *a* over all reads in the input data.

We also note that alternative uses for the quality scores during seed selection exist, depending on the nature of the overlap detection task. For example, given an algorithm that compares a small set of ‘query’ reads against a larger set of ‘index’ reads. This involves a different set of assumptions to use with Equation 1 which we will discuss as an alternative seed selection procedure at the end of this section.

### 2.2 Reverse-complement to forward frequency

Not all base-callers provide quality scores. In such cases the selection strategy must be determined using only seed frequency information.

If a sequencing technology or base-caller generates errors that are biased by the base content or context, this may be sufficient to create an improved selection strategy. Assuming that forward and reverse-complement reads of a given subsequence appear in equal proportions in the data, the ratio of the frequency of correct reads of the seed to the frequency of correct reads of its reverse complement should be 1:1. However if the type of errors are biased by base content then any given erroneous version of the seed should appear at different frequences from its reverse-complement. These unbalanced k-mers can then be assigned lower values, though this does rely on the bias in the errors to be greater than the difference in overall accuracy between the correct forward and reverse k-mers.

Of course, a seed containing an error may be equivalent to another seed that exists elsewhere in the data. This is a particular problem with low *k*, so we expect the ratio of forward to reverse-complement frequencies to become more useful as seeds increase in length. Algorithms that make use of this could also enforce constraints on the selected seeds to ensure dissimilarity, for example keeping a minimal edit distance between all pairs.

The strategy we use here calculates the balance between forward and reverse-complement as

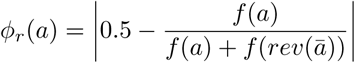
where *f*(*x*) is the frequency of subsequence *x* in the data, *x̅* is the complement of *x* and *rev*(*x*) its reverse.

### 2.3 Learning from external data

Another alternative to using a base-caller’s estimate of the probability of errors is to directly calculate the probability for each seed from existing data mapped to a known reference. A simple strategy *ϕ_t_*(*a*) is to use the proportion of these training reads’ subsequences that correctly map to a subsequence *a* in the reference. More sophisticated strategies could extend the domain of *ϕ* to include context around *a* in addition to its content alone.

In the evaluation section we shall test both training data drawn from the same contexts and drawn from data mapped to an entirely different genome from the input to simulate both selection strategies.

### 2.4 Local quality scores

Here we return to Equation 1 where we assumed *P*(*c*(*a, S*_0_)) = *P*(*c*(*a, S*_1_)). Instead we can use the local quality score information to make a better estimate of *P*(*c*(*a, S*_0_)), but continue to use the global mean quality for the probability of being correct in other sequences:

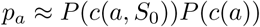
Thus the selection strategy is now dependent on the read *S*_0_ containing the potential seeds as well as the k-mer contents. In this paper we base selection on the product of the local quality score (at base offset by *k/*2 in *S*_0_) and the global quality score for the k-mer as described in Section 2.1. We will also consider using *ϕ_t_* and *ϕ_r_* in place of the global mean quality values.

## 3 Experimental setup

We determined per-seed values over all k-mers for the three selection strategies defined in the previous section, in addition to a lexicographic ordering to be used as a baseline. Values were calculated for *k* from 10 to 16.

Ground truth was determined by aligning reads to the reference using minialign[10] version 0.5.3. K-mers from reads that were contained in a ‘match’ alignment to the reference and contained no mis-matches were considered correct. All other k-mers in reads were considered incorrect. The fraction of correct instances for each k-mer were used as the ‘true’ or ideal selection strategy *ϕ_truth_*. K-mers that do not appear in any reads were assigned value 0.

### 3.1 Data

The input data used was a 60x coverage of E.Coli sequenced from an Oxford Nanopore MinION R9.4 and base-called with the albacore base-caller which provides quality scores. For training data with seeds in a similar context we used 200x coverage (900MB) of E.Coli from an Oxford Nanopore MinION R9.2 released by the Loman Labs, University of Birmingham^1^. For training data with unrelated seed context we used 30x coverage (1.9GB) of human chromosome 20 released by Oxford Nanopore Technologies^2^.

### 3.2 Evaluating selection strategies

To evaluate the quality of a selection strategy *ϕ* we compare the ranks of *ϕ*(*a*) to *ϕ_truth_*(*a*) for all k-mers *a* ∈ Σ^*k*^ that appear at least 3 times in the input data. The relationship is visualised as a heatmap with *ϕ_truth_* ranks on the y axis. Each row was normalised before plotting as there are disparities in the number of k-mers at each rank, particularly those with rank 0. In addition to the heatmap, the correlation coefficient of the ranks is calculated with higher values indicating a closer ordering of potential seeds to the ideal selection strategy.

We also consider the effect of seed filtering as in [2]. This is achieved by repeating the above procedure, but treating any k-mers not present in the reference as though they were not present in any reads. This is the best-case for seed filtering where all short reads are 100% accurate. In terms of the heatmaps, this just removes most data points from the bottom (rank 0) row. For the evaluation we therefore consider the effect the filtering has on the correlation of the remaining k-mers.

### 3.3 Evaluating local quality scores

Finally, we evaluate a local quality score metric for seed selection as described in Section 2.4. This is not a selection strategy, requires a specific (query-based) implementation, and therefore cannot be evaluated as correlation between ranks. For the ground truth we used all (top 1000) mappings found by minimap2 [**?**] to the E.Coli reference. The true overlap set at any given reference position is the set of all sequences with at least one mapping that spans that position.

We compared the utility of various seed selections at the task of finding all overlap sets based on a limited set of query sequences. The queries are taken from the first and last kilobase of each read, or whole reads if their length is below 2 kilobases. Query sequences were selected in batches and were used to select a global set of seeds, ensuring each query contains a minimum number of non-overlapping seeds and with the highest values present. Both query and index sequences are represented by all selected seeds (i.e. without additional sparcity constraints). Overlaps are then detected when the sets of seeds contain at least 3 matches that can be correctly ordered and are in approximately equal spacing. This is a fairly generic seed-based overlap detection algorithm with source code is available at https://github.com/jteutenberg/downpore if further details are required.

Under the assumption that these queries will be used for downstream assembly or consensus, the evaluation compares precision and recall of high quality overlap sets. We take a ‘high quality’ overlap set to be one in which at least 90% of all sequences matched by its query map to a single shared reference position. The overlap detection parameters were set to what were felt to be reasonable values at each *k* from 10 to 14 with the number of minimum seeds per query increasing from 15 to 45 over this range.

## 4 Results

Figure 1 shows the relationship between the true k-mer accuracy in the input data and that estimated by various selection strategies over a small selection of *k* values. A perfect correlation would appear as a dark diagonal line from lower left to upper right, showing that the selection strategy ranked k-mers in the same order as their actual rate of correctness in the data.

**Figure 1:**
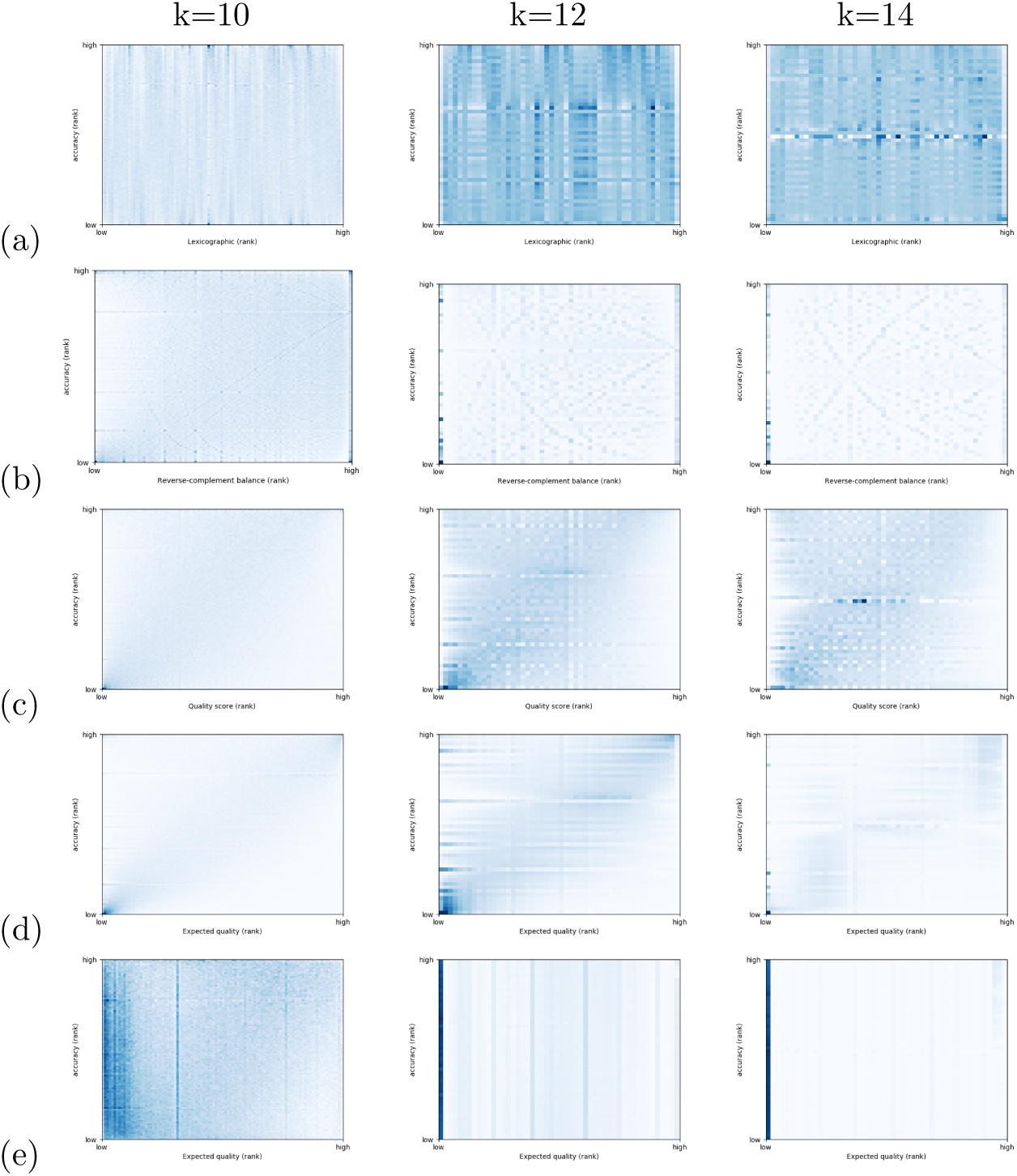
Examples of k-mer accuracy against (a) lexicographic ranks, (b) reverse-complement balance ranks, (c) base-caller quality score ranks, (d) same-genome trained accuracy ranks and (e) other-genome trained accuracy ranks for E.Coli data.

The top row (a) shows the lexicographic ordering. Some patterns do exist as k-mers containing the same motif on their left-hand side are nearby in the heatmap. These patterns suggests that k-mer quality is at least partially dependent on their subsequences, which in turn implies that global, per-k-mer measures of quality are useful for estimating the quality of any given subsequence.

The visually striking bands occur where many k-mers have small sample size in the data. This leads to ranks at fractions of the sample size containing many k-mers. For example, when only 3 examples of a k-mer exist in the data, its frequency (on the y axis) can only be at 0, 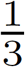, 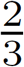 or 1. This situation occurs more frequently as *k* increases.

The second row (b) shows the rank associated with *ϕ_r_* – the ratio of frequency of each k-mer with its reverse-complement – with high ranks assigned to k-mers with balanced frequencies. While we expected this metric to become more informative as *k* increases and fewer single-base modifications overlap other correct k-mers. However, it appears that past *k* = 12 the low sample sizes lead to sensitive ratios between the frequency of k-mers and their reverse-complements and any useful correlation is lost. At *k* = 12 we see that a large number of low ranks of the ratios are assigned to few k-mers, and that most k-mers appear balanced irrespective of their actual accuracy.

Three further comparisons (c) - (e) were made that rely on additional data: quality scores from the base-caller; known k-mer accuracies from other data on the same genome; and known k-mer accuracies from a different genome.

Both the quality scores and the k-mers from the same genome show a linear correlation for all *k* tested. The quality score heatmaps show an increased concentration of k-mers above the diagonal – a population in which k-mer ranks have been underestimated.

Using data from the human chromosome 20 appears to have two populations of k-mers: those that correlate with true k-mer ranks (only visible when *k* = 10); and those assigned low ranks independent of their true rank. It is understandable that as *k* increases the number of k-mers shared between the different genomes will reduce, with the left-most column representing k-mers that do not appear in the training reference but do appear in the input data’s reference.

Figure 2 gives the correlation coefficient between the ranks of all selection strategeies, the lower *k* values of which correspond to those shown in Figure 1. As expected, the lexicographic ordering is uncorrelated at all *k* and the ranks taken from human chromosome 20 are no better for *k >* 10.

**Figure 2:**
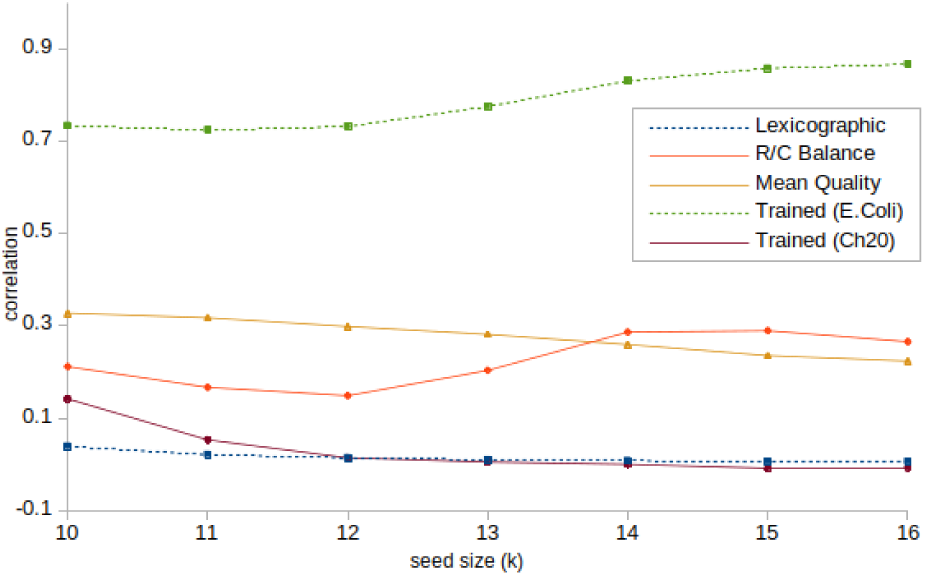
Correlation coefficients for the E.Coli seed metrics to accuracy data for *k* up to 16, including data shown in Figure 1.

Of the remainder, using data trained on the same genome so the k-mers appear in the same contexts is the best of these selection strategies. However, the more accessible strategies based on mean quality scores and the reverse-complement ot forward ratio also provide reasonable correlation with the best ordering of k-mers.

### 4.1 Combining with accurate seeds

Figure 3 repeats the experiment shown in Figure 2 but ignoring all seeds not present in the reference. It has been shown that removing known irrelevant seeds improves overlap detection [2], we wish to determine whether our proposed selection strategies can estimate the accuracy of the remaining k-mers.

**Figure 3:**
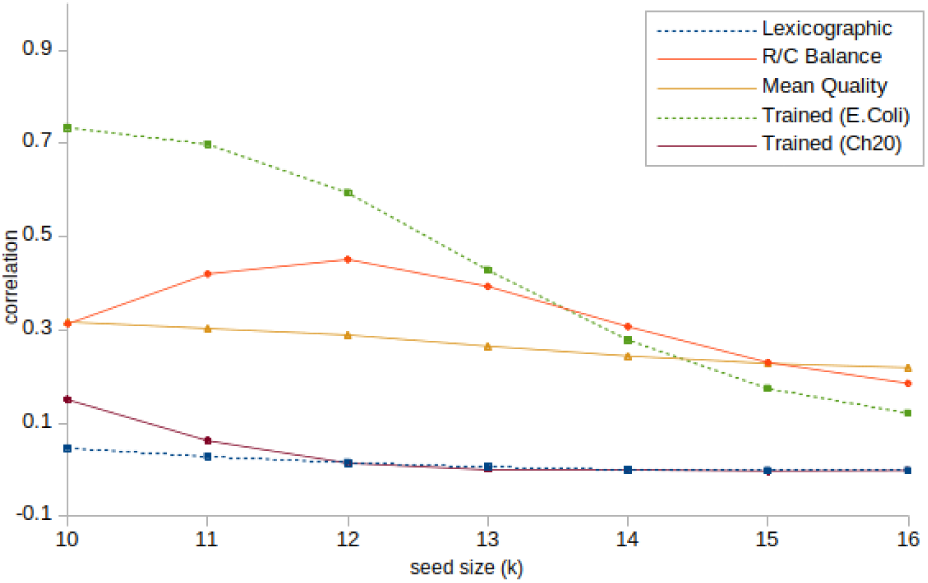
Correlation coefficients for the data in Table 1, restricted to k-mers present in the reference.

The two low-correlation strategies were unchanged. The mean quality scores were also correlated to the same extent on the filtered set of k-mers, so this strategy continues to provide useful information beyond just indicating k-mers that are always incorrect.

The values trained on other E.Coli data become less relevant as *k* increases. This indicates that there is substantial overlap between the information provided by the training data and the filtering process.

Finally, an interesting increase in correlation between *ϕ_r_* for *k* below 14 was seen. In terms of the heatmaps in Figure 1 the filtering removes most of the bottom row (k-mers with no correct instances in the reads), including the large number of k-mers at the origin. Particularly in the *k* = 10 figure there appears to be more k-mers present at higher ranks than lower, thus once the values at the origin are removed a better linear relationship shifted to the right may be found. By *k* = 14, visually this is no longer apparent which fits with the return to similar correlation of about 0.3.

### 4.2 Effect on overlap detection

The results of determining overlap sets from batched 1kB queries is shown in Figure 4. The parameters used appear to be more suited to the low *k* values with both precision and recall dropping off rapidly as it increases. The *k* = 14 values are not shown for readability, but the same trend seen in the figure continues.

**Figure 4:**
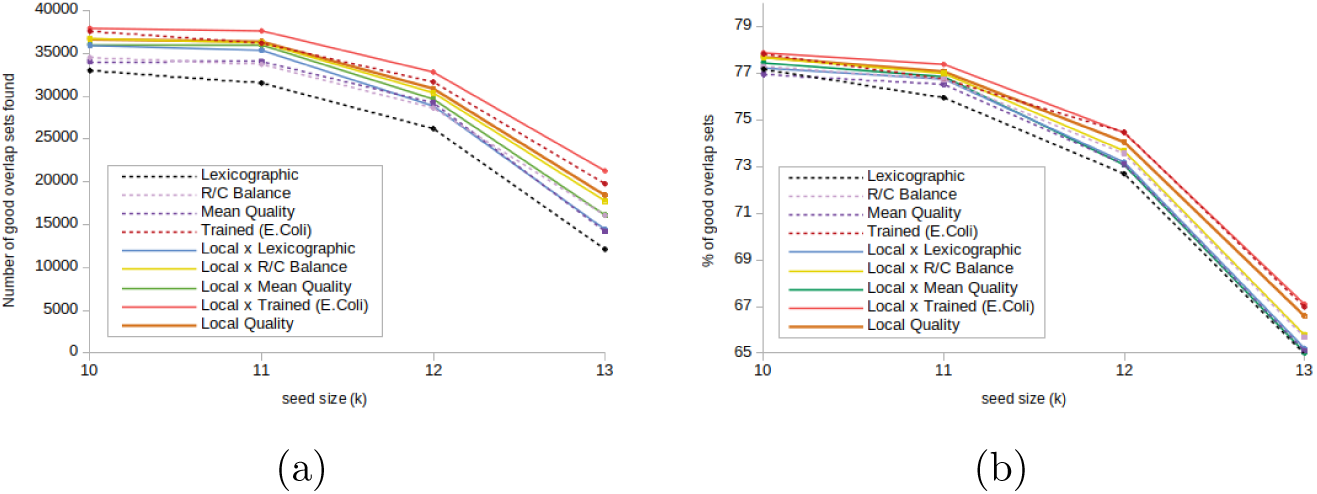
Effect of seed selection on overlap set quality at various *k*, as (a) recall: number of overlaps (in thousands) found with at least 90% component sequences correctly identified and (b) precision: percentage of all overlap sets with at least 90% component sequences correctly identified.

Firstly, the inclusion of local quality scores always improves both precision and recall of an existing selection strategy. This includes the best-performing strategy based on k-mer accuracies taken from the same E.Coli source.

Secondly, asside from the mean quality scores *ϕ_q_* at *k* = 10 (which performed unusually poorly), the ordering of selection strategies was unchanged over *k*. In addition, the ordering of selection strategy by recall is the same as by precision– so we have strictly better selection strategies for this task.

Finally, using local quality values alone actually performs better than combining with global measures *ϕ_r_* or *ϕ_q_*. This is unexpected given that we have shown these global measures do correlate with the expected correctness of a seed in other sequences.

Overall, replacing a lexicographic ordering with a strategy that includes local quality scores improved recall by 10-15% while marginally increasing precision.

## 5 Conclusion

We have shown that uninformed lexicographic ordering of k-mers can be improved on using simple per k-mer metrics. Examples that produce k-mer rankings that correlate with their probability of being correct in reads have been been presented based on k-mer frequency alone (e.g. a reverse-complement to forward ratio); based on base-callers’ quality scores; or based on external training data. During overlap detection there is no excuse for comparing potential seeds based on their lexicographic order.

We have also provided an improved seed selection scheme for applications that determine overlaps based on a small query set. Using a base-caller’s quality scores on the specific position of a potential seed in a query further improves on the global per k-mer metrics. In addition, all seed selection strategies here have been shown to improve an example overlap detection algorithm in line with their correlation to k-mer correctness.

From those strategies considered in this paper we recommend the replacement of lexicographic ordering by mean quality scores per k-mer. This correlates well with the likelihood of a seed being shared by overlapping sequences, is easily obtained, and is stable across a range of *k* values. For those applications that permit it, the local quality score of a query seed should always be combined with this value.

## Footnotes

1 https://s3.climb.ac.uk/nanopore/E_coli_K12_1D_R9.2_SpotON_2.pass.fasta

2 https://github.com/nanoporetech/ONT-HG1/blob/master/CONTENTS.md

